# Pericytes and Wnt signaling induce functional blood-brain barrier phenotype in human iPSC-based model

**DOI:** 10.1101/2025.09.26.678752

**Authors:** Henrique Nogueira Pinto, Nine R. Kok, Philipp C. Hauger, Manon Karsten-van Diepen, Michael de Kok, Nanne J. Paauw, Susanne M.A. van der Pol, Joline P. Nugteren-Boogaard, Peter L. Hordijk, Stephanie D. Beekhuis-Hoekstra, Susan Gibbs, Nienke M. de Wit, Helga E. de Vries

## Abstract

The blood-brain barrier (BBB), formed by brain microvascular endothelial cells (BMECs), restricts vascular permeability through tight junctions, selective transporters, and low transcytosis. BBB dysfunction contributes to cerebrovascular and neurodegenerative disease, yet current human in vitro models recapitulate only a subset of BMEC features. Here, we describe a strategy generate BMECs (hiBMECs) from human induced pluripotent stem cell-derived endothelial cells by co-culture with isogenic brain pericytes and activation of Wnt/β-catenin signaling. The resulting hiBMECs display barrier properties, active efflux transporters, and appropriate inflammatory responses. Transcriptomic profiling revealed convergence of pericyte-derived cues and Wnt/β-catenin activation on ETS1, SMAD3/4, and PPARγ transcriptional networks, establishing a gene signature closely matching the adult human BBB. Downstream analysis revealed that hiBPC cues engaged sphingosine-1-phosphate, TGF-β, and angiopoietin/Tie2 pathways, which were further regulated by canonical Wnt activation. These findings uncover a synergistic mechanism by which brain pericytes and Wnt/β-catenin signaling orchestrate BMEC differentiation and function, providing mechanistic insight into human BBB development and an improved hiPSC-derived BBB model for future drug screening and disease modeling.

**Graphical Abstract:** 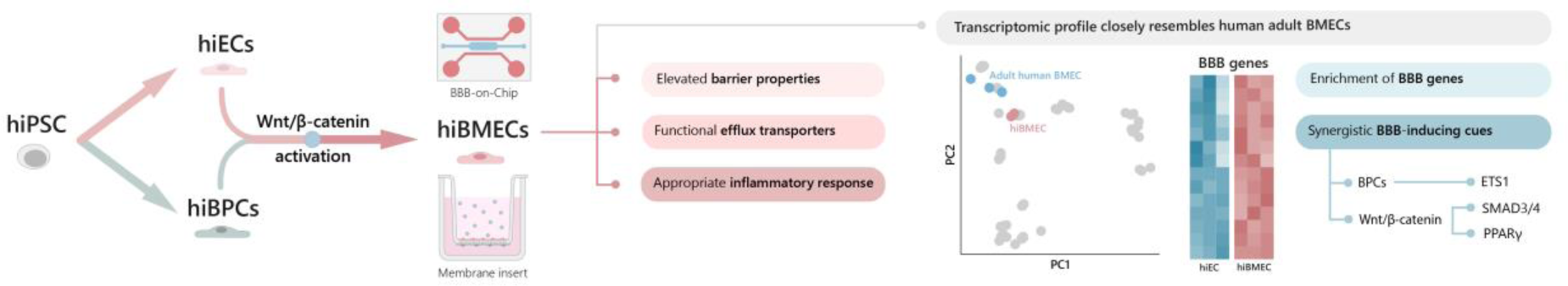

**The paper explained:** *PROBLEM:* The blood–brain barrier (BBB) protects the brain by tightly regulating the passage of molecules and cells. Its dysfunction contributes to disorders such as stroke, dementia, and multiple sclerosis. Yet, existing human in vitro models fail to capture the full complexity of BBB biology, limiting our ability to study disease mechanisms or test brain-targeted drugs.

*RESULTS:* We discovered that two signals are essential for generating functional human BBB endothelial cells from stem cells: cues from brain pericytes and activation of the Wnt/β-catenin pathway. Together, these signals enabled endothelial cells to form tight barriers, operate transporters, and mount appropriate responses to inflammation. Transcriptomic analyses of the resulting cells revealed a gene signature closely matching the adult human BBB and identified how pericyte- and Wnt-activated pathways converge on specific transcriptional programs driving BBB identity.

*IMPACT:* This study provides both a molecular framework for in vitro BBB development and a reliable and reproducible human BBB model. This platform can be applied to explore BBB dysfunction in neurological disease and to accelerate the development of drugs that need to reach the brain or target the brain vasculature.

## Introduction

The brain is protected from blood-borne insults by the blood-brain barrier (BBB), formed by a continuous layer of brain microvascular endothelial cells (BMECs), which are supported in their function by pericytes, astrocytic endfeet, and basement membranes (1). BMECs are specialized endothelial cells (ECs) with properties that differ from ECs outside the central nervous system. BMECs establish tight junctions (TJs), present low levels of vesicular traffic and do not display fenestrations, which leads to a reduced vascular permeability. Influx and efflux transporters on their membranes regulate the traffic of biomolecules between the blood and the brain. BMECs are also responsive to inflammation by upregulating cell adhesion molecules (CAMs), pro-inflammatory cytokines and proteins involved in antigen presentation (2–5).

BBB dysfunction is an early hallmark of multiple neurodegenerative and neuroinflammatory diseases, such as Alzheimer’s disease (6–8), Parkinson’s disease (9), and multiple sclerosis (10), making it an interesting therapeutic target due to its critical role in disease initiation and progression. By enhancing barrier integrity, modulating immune cell trafficking, or improving drug delivery to the brain, BBB-targeted therapies may help mitigate early pathological changes such as increased permeability, impaired amyloid-β clearance, and neuroinflammation, therefore preventing or slowing neuronal damage and cognitive decline (11). Moreover, efflux transporters play a major role in drug resistance at the BBB by restricting the entry of xenobiotic compounds into the brain, limiting treatment of brain disorders (1). Therefore, there is a need for a reliable tool to investigate the molecular mechanisms underlying human BBB pathophysiology and to screen drugs and drug delivery systems that are able to cross and/or repair the BBB.

*In vivo* models are important tools to study systemic responses in a complex environment. However, beyond the ethical concerns surrounding the use of animals in biomedical research, there is also compelling evidence pointing to significant differences between animals and humans, particularly in BBB-related gene expression, signaling pathways, protein homology, and brain vascular architecture (12). These limitations have driven the development of *in vitro* systems that utilize human-derived materials to more accurately model the human BBB. Despite their relevance, primary human BMECs are hard to obtain and to expand in vitro and tend to lose BBB characteristics over time (13). In contrast, immortalized BMEC lines can be readily expanded but have limited transendothelial electrical resistance (TEER) and TJ development (14). CD34^+^ hematopoietic stem cells (HSC) isolated from umbilical cord blood, can obtain BMEC-like properties when co-cultured with bovine brain pericytes, which can be further enhanced with a combination of small molecules (15, 16); however, donor-to-donor variability, scalability, and the use of non-human pericytes might compromise the translatability of the model.

To overcome these limitations, multiple protocols to generate BMEC-like cells from human induced pluripotent stem cells (hiPSCs) have been developed. HiPSCs can be obtained from multiple donors, expanded in large scale, and differentiated into multiple cell types, which enables the development of human isogenic *in vitro* platforms for disease modelling or drug screening (17). Researchers have successfully generated BMEC-like cells with high TEER, low permeability, and active efflux transporters, using unconditioned (UMM-BMECs) (18) and chemically defined (DMM-BMECs) (19) medium methods. However, these cells have limited response to inflammatory stimuli and co-express epithelial markers, which are the main drivers of the elevated barrier properties (20, 21). Other methods can generate BMEC-like cells that respond to inflammatory stimuli but present relatively low TEER values and do not display efflux transporter activity (20, 22, 23). These limitations illustrate the need for an improved method to generate BMECs encompassing the key features of the BBB.

In development, blood vessels from the perineural vascular plexus (PNVP) invade the neural tube following a gradient of growth factors (24). The ECs from the PNVP do not possess BBB properties. Neural crest-derived pericytes are recruited to the invading ECs, stabilizing nascent vessels and inducing barriergenesis (25, 26). Concomitantly, neural progenitor cells secrete Wnt ligands that further enhance angiogenesis and BMEC maturation via the Wnt/β-catenin signaling pathway (27–29). Therefore, the presence of pericytes and the activation of Wnt, together, are essential to induce the BMEC phenotype in ECs.

Inspired by BBB development *in vivo*, we established a method to differentiate hiPSCs into BMECs (hiBMECs) with mature TJs, active efflux transporters, and a fully endothelial phenotype by co-culturing hiPSC-derived ECs (hiECs) with hiPSC-derived brain pericytes (hiBPCs) and activating the Wnt/β-catenin pathway. Using this approach, we have characterized the synergistic effects of hiBPCs and Wnt signaling on human *in vitro* BMEC lineage commitment from naïve endothelium and established an enhanced hiPSC-based BBB model that is amenable to disease modeling and drug screening.

## Results

### BMEC identity is induced by pericytes and Wnt/β-catenin signaling in hiECs

To generate hiBMECs, we differentiated endothelial cells from hiPSCs through mesodermal lineage commitment and vascular specification, by adapting previously described protocols (30, 31) (Fig 1A). We purified the resulting platelet endothelial cell adhesion molecule (PECAM-1)^+^ cells and co-cultured them in a porous membrane insert with isogenic neural crest-derived hiBPCs that were differentiated in parallel (Fig EV1A). On the final two days, cells were stimulated with the glycogen synthase kinase 3 (GSK3) inhibitor CHIR, activating the Wnt/β-catenin pathway.

**Figure 1.**
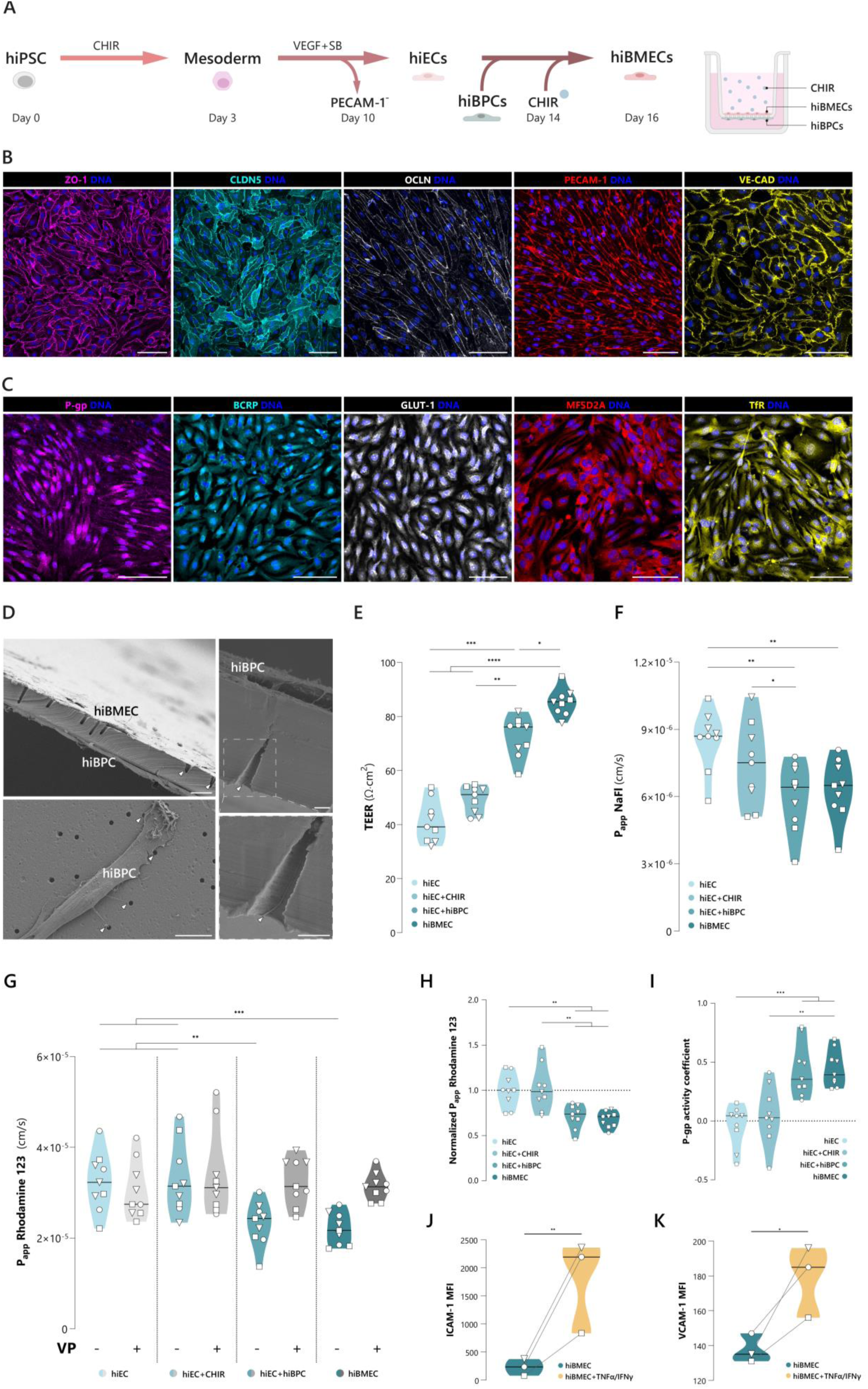
Differentiation of hiBMECs from hiECs through hiBPC co-culture and Wnt/β-catenin activation. **(A)** Overview of the hiBMEC differentiation protocol and schematic illustration of the experimental set-up. **(B)** Expression of tight junction proteins ZO-1, CLDN5 and OCLN, and endothelial markers PECAM-1 and VE-CAD in hiBMECs using immunocytochemistry. Scale bars: 100 μm. **(C)** Expression of efflux transporters P-gp and BCRP, and other BBB-enriched transporters GLUT-1, MFSD2A and TfR in hiBMECs using immunocytochemistry. Scale bars: 100 μm. **(D)** Scanning electron microscopy of hiBMECs and hiBPCs co-cultured on a membrane insert. Arrows indicate cellular processes protruding into and across the pores. Scale bars: 10 μm (left); 2 μm (right). **(E)** Transendothelial electrical resistance (TEER in Ω·cm_2_) of monolayers of hiEC, hiEC+CHIR, hiEC+hiBPC and hiBMEC. **(F)** Apparent permeability (P_app_) to sodium fluorescein (NaFl) of hiEC, hiEC+CHIR, hiEC+hiBPC and hiBMEC. **(G)** P_app_ to P-gp substrate Rhodamine 123 (Rho123) of hiEC, hiEC+CHIR, hiEC+hiBPC and hiBMEC, with or without treatment with P-gp inhibitor verapamil (VP). **(H)** Rho123 P_app_ of untreated hiEC+CHIR, hiEC+hiBPC and hiBMEC, normalized to hiEC. **(I)** P-gp activity coefficient of hiEC, hiEC+CHIR, hiEC+hiBPC and hiBMEC, based on the ratio of Rho123 P_app_ between VP-treated and untreated cells. **(J-K)** Flow cytometry analysis of mean fluorescent intensity (MFI) of ICAM-1 and VCAM-1 in hiBMECs with and without TNFα and IFNγ stimulation (25 IU/mL, 16 hours). **(E-I)** Points represent the average of 3 technical replicates from 3 independent differentiations and 3 independent experiments. Ordinary one-way ANOVA. \*\*\*\**P* < 0.001, \*\*\**P* < 0.005, \*\**P* < 0.01, \**P* < 0.05. **(J-K)** Points represent the average of 3 technical replicates from 3 independent differentiations. Two-tailed paired student t-test. \*\**P* < 0.01, \**P* < 0.05. **(E-K)** Symbols represent the different hiPSC lines: ▽ = GB2a, ◯ = GB4d, □ = TMO.

HiBMECs grew into a confluent monolayer and displayed a continuous expression, at the cell-cell contacts, of the tight junction proteins zonula occludens-1 (ZO-1), claudin-5 (CLDN5), and occludin (OCLN), the cell adhesion molecule PECAM-1, and the adherens junction protein vascular endothelial cadherin (VE-CAD) (Fig 1B). HiBMECs expressed the efflux transporters P-glycoprotein (P-gp) and breast cancer resistance protein (BCRP), as well as glucose transporter 1 (GLUT1), major facilitator superfamily domain-containing protein 2 (MFSD2A) and transferrin receptor (TfR), all of which are enriched in human BMECs in comparison to peripheral ECs (Fig 1C). Additionally, we performed scanning electron microscopy for a detailed visualization of the model and observed that hiBPCs extended cellular processes through the membrane pores, directly contacting the basal side of hiBMECs (Fig 1D).

To assess the barrier properties of hiBMECs, we compared the TEER of hiBMECs, hiECs, hiECs stimulated with CHIR (hiEC+CHIR) and hiECs co-cultured with hiBPCs (hiEC+hiBPC). HiECs displayed a basal TEER of ∼40 ohm·cm^2^. The addition of CHIR led to a slight increase to ∼50 ohm·cm^2^ of average TEER, albeit not significant. The co-culture with hiBPCs induced a significant increase of TEER to ∼75 ohm·cm^2^. Notably, the combination of hiBPC co-culture and CHIR stimulation led to a TEER of over 80 ohm·cm^2^ in hiBMECs (Fig 1E). To assess paracellular diffusion, we determined the apparent permeability (P_app_) of hiBMECs to sodium fluorescein (NaFl) as an additional benchmark for BBB function. In line with the TEER results, the addition of CHIR to hiECs did not lead to a significant decrease of P_app_. The co-culture with hiBPCs, however, significantly decreased the NaFl P_app_ to ∼6×10^−6^ cm/s, independently of CHIR stimulation (Fig 1F).

We next determined P-gp activity in hiBMECs, using Rho_123_ as P-gp substrate and verapamil (VP) as P-gp inhibitor. No P-gp activity was observed for hiECs and hiECs+CHIR, as determined by the unchanged Rho_123_ P_app_ upon VP treatment and consequent null P-gp activity coefficient (Fig 1G-I). Conversely, the Rho_123_ P_app_ in hiBMECs and hiECs+hiBPCs decreased in comparison with hiECs, and increased upon VP treatment, leading to a positive P-gp activity coefficient and indicating P-gp function in these conditions. Additionally, we assessed the effect of activating pathways implicated in BBB function and P-gp induction using soluble factors [cyclic adenosine monophosphate + CHIR + A-83-01 (15), retinoic acid (32), GW3965 (33, 34), RepSox (35), forskolin (36), epidermal growth factor (37), vitamin D3 (38), dexamethasone (39), rifampicin (40), and valproic acid (41)], but none of the tested soluble factors induced P-gp activity in hiECs (Appendix Fig S1). Similarly, hiEC that were co-cultured with hiPSC-derived astrocytes, or stimulated with astrocyte or pericyte conditioned medium for 7 days did not show P-gp activity.

Finally, we verified the capacity of differentiated hiBMECs to respond to inflammatory stimuli. To do so, we stimulated hiBMECs with pro-inflammatory cytokines TNFα and IFNγ and performed flow cytometry for intercellular adhesion molecule 1 (ICAM-1) and vascular cell adhesion molecule 1 (VCAM-1). When comparing inflamed to unstimulated hiBMECs, we observed a significant increase in ICAM-1 (8.7 times) and VCAM-1 (1.3 times) expression (Fig 1J-K, Appendix Fig S2A-B) and in VCAM-1^+^ and ICAM-1^+^ cell populations (Fig EV2A-B, Appendix Fig S2C-D), demonstrating an appropriate endothelial response to inflammation without compromising the monolayer integrity.

### HiBMECs can be differentiated in a self-organized 3D microvascular model

In order to mimic physical aspects of the BBB microenvironment, we aimed to differentiate hiBMECs in a microfluidic system that integrates three-dimensionality and flow. Our BBB-on-Chip (BBBoC) was developed using the AIM Biotech idenTx 9 platform, in which hiECs and hiBPCs were embedded in a fibrin gel and co-cultured in the central channel of the chips (see workflow in Fig 2A). Over seven days, a self-assembled 3D vascular network was formed (Fig 2B). These networks were lumenized and perfusable, with an average vessel diameter of 60 μm, average branch length of 110 μm, total network length of 20 mm and 200 branching points per network (Fig 2C-D, Fig EV2A-C).

**Figure 2.**
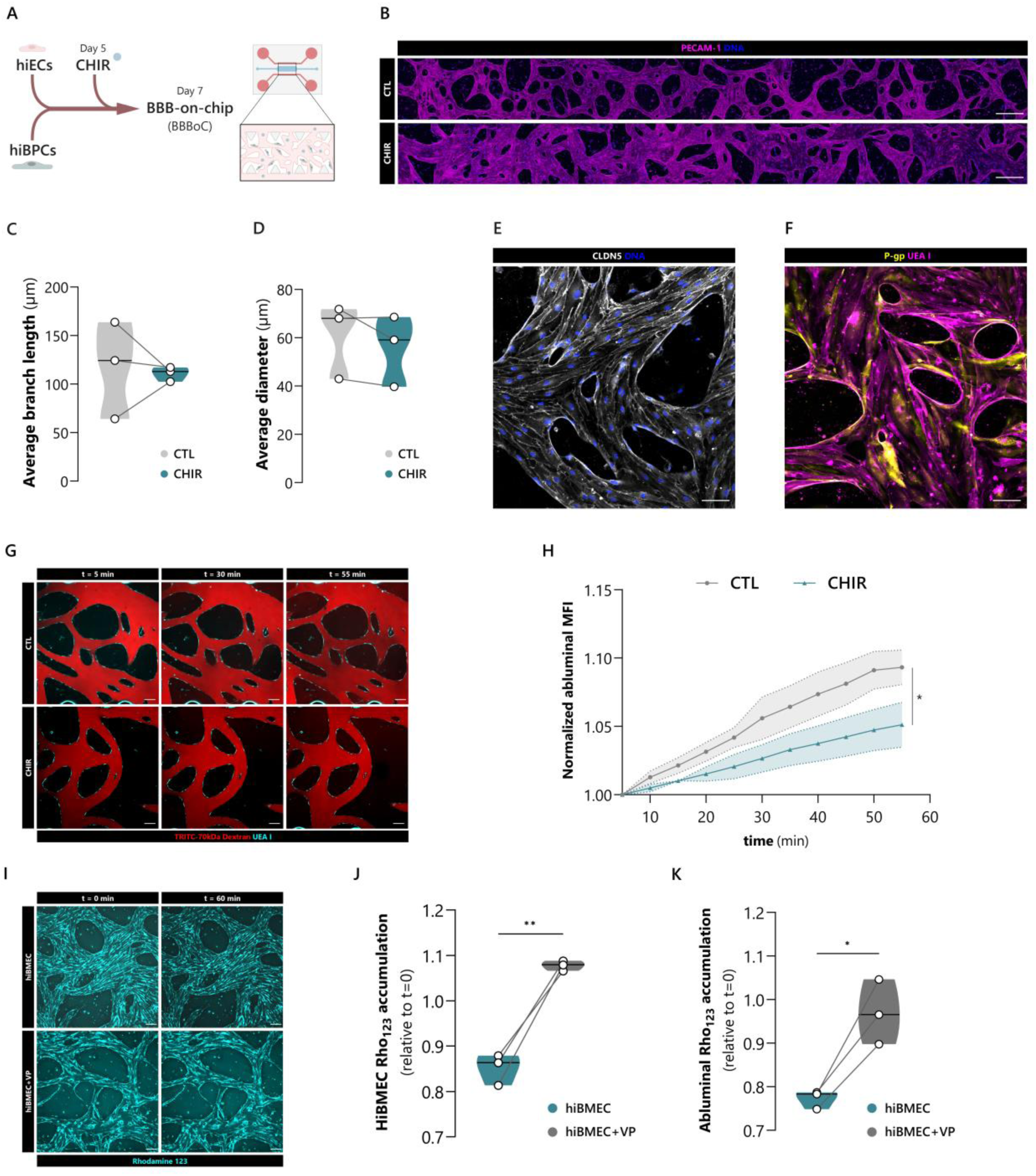
Differentiation of hiBMECs in a self-organized 3D microvascular model. **(A)** Overview of the BBB-on-chip (BBBoC) protocol and schematic illustration of the experimental set-up. **(B)** Representative immunofluorescence image of a microvascular network formed by hiBMECs and hiBPCs. Vessels are stained with PECAM-1 and span across the entire microfluidic channel. Top image shows a maximum intensity projection of the Z stack and bottom image shows a single Z slice, highlighting the vessel lumens. Scale bars: 500 μm. **(C)** Average vessel diameter and **(D)** average branch length of BBBoCs treated with CHIR at day 5 (CHIR) or untreated (CTL), in μm. Points represent 3 BBBoCs from 3 independent differentiations. **(E)** Immunocytochemistry of tight junction protein CLDN5 and **(F)** efflux transporter P-gp, present in 3D hiBMECs. Scale bar: 100 μm. **(G)** Representative images of 3 timepoints (5, 30 and 55 minutes) of permeability assay using TRITC-labeled 70kDa dextran in BBBoCs with or without the addition of CHIR for BMEC commitment. Vessels are stained with UEA I. Scale bars: 100 μm. **(H)** Quantification of abluminal main fluorescence intensity of TRITC-labeled 70kDa dextran in the gel surrounding the vessels over 55 minutes. Values were normalized to fluorescence at 5 minutes. Mean ± SD, Two-way repeated measures ANOVA with Greenhouse-Geisser correction, *P < 0.05, n = 3. **(I)** Representative images of 2 timepoints of P-gp activity assay in BBBoCs using Rhodamine 123 (Rho_123_) as P-gp substrate and verapamil (VP) as P-gp inhibitor. Scale bars: 100 μm. **(J)** Quantification of Rho_123_ accumulation in hiBMECs and **(K)** abluminally, normalized to 0 hours. Points represent the average of 3 technical replicates from 3 independent experiments. Two-tailed paired student t-test. \*\**P* < 0.01, \**P* < 0.05.

We observed using immunocytochemistry that vessels were positive for CLDN5 and P-gp (Fig 2E-F), and in close contact with hiBPCs (Fig EV2D). CHIR treatment for 48 hours decreased the permeability of the vascular network to 70kDa TRITC-dextran (Fig 2G-H), without affecting average vessel diameter, average branch length, total network length, and number of branching points (Fig 2C-D, Fig EV2B-C). To further confirm the BMEC identity, we performed the P-gp functional assay on the BBBoC and observed that the Rho_123_ fluorescence intensity in hiBMECs and on the abluminal side was increased in VP-treated chips (Fig 2I-K), indicating P-gp activity. Together, this indicates that hiECs were successfully differentiated into hiBMECs on-chip over seven days of co-culture with hiBPCs and 48 hours of CHIR stimulation.

### Transcriptomic profile of hiBMECs closely resembles adult human BMECs

To dissect the transcriptomic profile of the distinct differentiated EC types, we performed bulk RNA sequencing on hiECs, hiECs+CHIR, hiECs+hiBPCs and hiBMECs. Differential gene expression analysis of hiBMECs and hiECs identified 2021 differentially expressed genes (DEGs, adjusted p-value below 0.05 and fold change above 1.5), of which 847 were significantly upregulated and 1258 were significantly downregulated in hiBMECs (Fig 3A).

**Figure 3.**
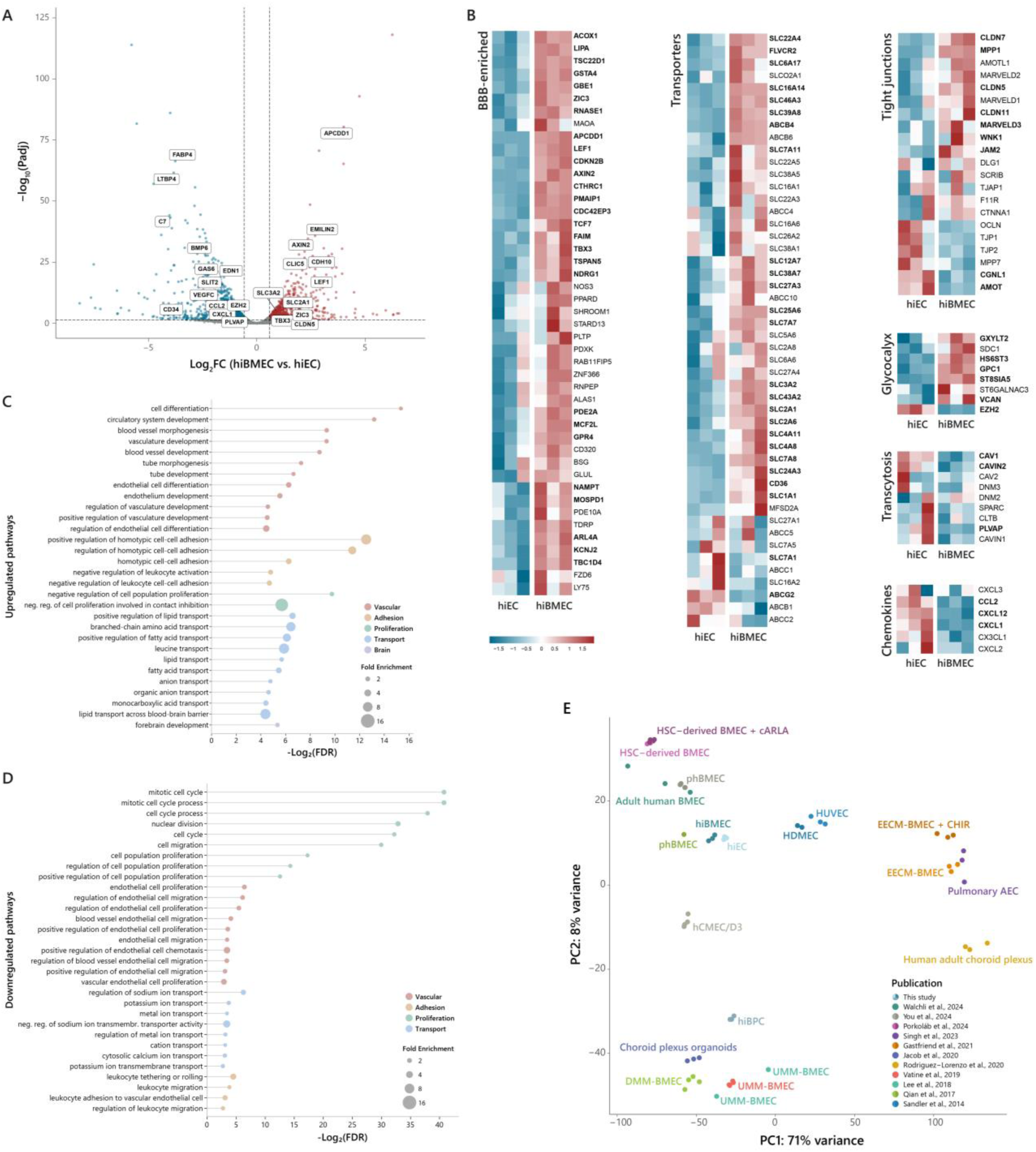
Transcriptomic characterization of hiBMECs. **(A)** Volcano plot showing differentially expressed genes of hiBMECs vs hiECs. Upregulated genes (adjusted *p* value < 0.05 and fold change > 1.5) are displayed in red and downregulated genes (adjusted *p* value < 0.05 and fold change < −1.5) are displayed in blue. A number of BBB-related transcripts are highlighted. N=3 per condition, GB4d hiPSC line. *P* values were determined using the Wald test and adjusted for multiple testing using the Benjamini-Hochberg procedure. **(B)** Z-score heatmaps of gene expression profiles across hiEC and hiBMEC. Selected transcripts are enriched in brain ECs vs. peripheral ECs and/or related with solute transport, tight junctions, glycocalyx, transcytosis and chemotaxis. Significantly changed genes are marked in bold. **(C)** Gene ontology enrichment analysis of upregulated and **(D)** downregulated gene sets in hiBMECs vs. hiECs. Gene Ontology Biological Process (GO:BP) terms are categorized into vascular, adhesion, proliferation, transport and brain, and ranked based on their false discovery rate (FDR). **(E)** Principal component analysis (PCA) plot illustrating relative relationship of 65 distinct cell samples across 12 library preparations from previously published and newly generated bulk RNA sequencing data, including hiECs and hiBMECs.

Amongst the induced transcripts, we found 32 genes that are enriched at the BBB *in vivo* (15, 42) (Table S1), such as the Wnt target genes *APCDD1*, *AXIN2*, *CTHRC1* and *LEF1*, and the transcription factors *TBX3*, *TCF7, TSC22D1* and *ZIC3*. Similarly, we observed a trend of increased expression of other BBB-enriched genes in hiBMECs, such as *BSG*, *PPARD*, *NOS3*, and *FZD6* (Fig 3B), although not significant. Moreover, we observed that some of the most downregulated genes were involved in barrier-disruptive processes like angiogenesis (*EDN1*, *VEGFC*, *SLIT2*) and inflammation (*FABP4*, *LTBP4*, *C7*, *GAS6*) (Fig 3A). Notably, the expression of key endothelial transcripts was significantly higher compared to epithelial transcripts, confirming the vascular endothelial identity of hiBMECs (Fig EV3).

Numerous solute carrier (SLC) transporters were upregulated in hiBMECs vs. hiECs, including GLUT-1 (*SLC2A1*), CD98hc (*SLC3A2*), EAAT3 (*SLC1A1*) and many others (Fig 3B). HiBMECs were also enriched in other BBB transporters, such as the choline and heme transporter *FLVCR2* and the fatty acid transporter *CD36*, and showed a trend towards increased expression of the BBB-specific transporter *MFSD2A*, although not significant. Conversely, the expression of ATP-binding cassette (ABC) transporters was not extensively altered in hiBMECs vs. hiECs, with only an upregulation of *ABCB4* and downregulation of *ABCG2*.

Regarding changes in genes related with TJs, we observed a significant enrichment of *CLDN5*, *7* and *11*, *AMOTL1*, *MARVELD3*, *WNK1*, *JAM2*, and *MPP1*, and a decreased expression of *AMOT* and *CGNL1* in hiBMECs compared with hiECs (Fig 3B). Additionally, we observed a downregulation of *EZH2*, a transcriptional repressor of various endothelial glycocalyx-associated genes, such as *GXYLT2* and *HS6ST3* (43). In agreement, these and other negatively charged glycocalyx-related genes were significantly upregulated in hiBMECs (Fig 3B).

HiBMECs displayed an overall downregulation of genes involved in transcytosis, such as *PLVAP*, *CAV1* and *CAVIN2*. Similarly, hiBMECs showed a decreased expression of genes encoding chemokines that stimulate the migration and activation of immune cells, including *CXCL12*, *CCL2*, and *CXCL1* (Fig 3B).

Gene ontology (GO) analysis of the upregulated DEGs in hiBMECs versus hiECs revealed an enrichment of pathways related to vasculature and forebrain development, endothelial differentiation, homotypic cell-cell adhesion, contact inhibition, and transport of lipids and amino acids (Fig 3C). Downregulated DEGs were involved in pathways related to cell cycle, proliferation, migration, ion transport, and leukocyte adhesion (Fig 3D).

To benchmark the transcriptomic profile of hiBMECs, we performed a principal component analysis (PCA) on our dataset together with previously published datasets of freshly isolated adult human BMECs (44), hiPSC-derived BMEC-like cells (15, 19, 23, 45, 46), primary and immortalized ECs lines (47–49), and choroid plexus explants and organoids (50, 51) (Fig 3E). BMEC-like cells derived from defined medium method (DMM-BMECs) and unconditioned medium method (UMM-BMECs) clustered with choroid plexus epithelial organoids, while BMEC-like cells derived from extended endothelial cell culture method (EECM-BMECs) were grouped with pulmonary arterial endothelial cells. Conversely, hiBMECs clustered closely to freshly isolated adult human BMECs and primary human BMECs (phBMECs), indicating that our hiBMECs share substantial transcriptional similarities with adult human BMECs.

### HiBMECs respond appropriately to inflammatory stimuli

To evaluate the response of hiBMECs to inflammation, we stimulated hiBMECs, hiECs, hiECs+CHIR and hiECs+hiBPCs with the pro-inflammatory cytokines TNFα and IFNγ (25 IU/mL) for 16 hours and performed bulk RNA sequencing on these cells. The PCA indicated profound transcriptomic changes induced by this stimulation in all conditions (Fig 4A).

**Figure 4.**
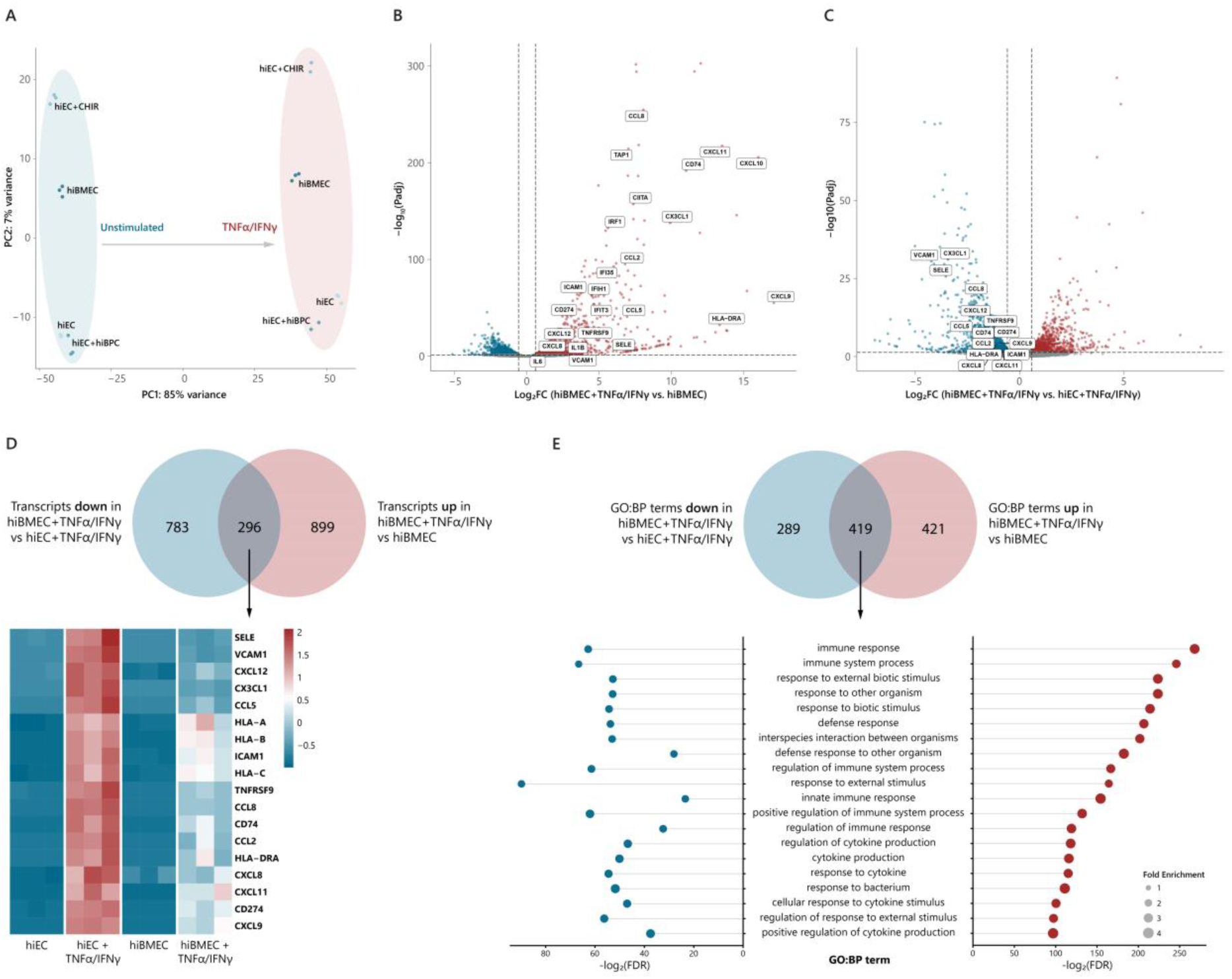
Inflammatory response of hiBMECs to TNFα and IFNγ stimulation. **(A)** PCA plot of hiEC, hiEC+CHIR, hiEC+hiBPC and hiBMEC with and without TNFα and IFNγ stimulation (25 IU/mL, 16 hours). **(B)** Volcano plot showing differentially expressed genes of hiBMEC+TNFα/IFNγ vs hiBMEC and **(C)** hiBMEC+TNFα/IFNγ vs hiEC+TNFα/IFNγ. Upregulated genes (adjusted *p* value < 0.05 and fold change > 1.5) are displayed in red and downregulated genes (adjusted *p* value < 0.05 and fold change < −1.5) are displayed in blue. Key inflammation-related transcripts are highlighted. N=3 per condition, GB4d hiPSC line. *P* values were determined using the Wald test and adjusted for multiple testing using the Benjamini-Hochberg procedure. **(D)** Venn diagram of transcripts upregulated in hiBMEC+TNFα/IFNγ vs hiBMEC and downregulated in hiBMEC+TNFα/IFNγ vs hiBMEC. The expression profile of key overlapping genes in hiEC, hiEC+TNFα/IFNγ, hiBMEC and hiBMEC+TNFα/IFNγ is shown in the Z-score heatmap below. **(E)** Venn diagram of GO:BP terms enriched in the gene sets upregulated in hiBMEC+TNFα/IFNγ vs hiBMEC and downregulated in hiBMEC+TNFα/IFNγ vs hiEC+TNFα/IFNγ. The 20 overlapping terms with lower FDR in hiBMEC+TNFα/IFNγ vs hiBMEC are highlighted below in a lollipop plot.

Differential gene expression analysis of inflamed and control hiBMECs identified 1195 upregulated and 798 downregulated DEGs (Fig 4B). Inflamed hiBMECs upregulated key endothelial cell adhesion molecules, such as *ICAM1* (13-fold), *VCAM1* (10-fold) and *SELE* (74-fold) (Fig EV4A). We also observed an upregulation of multiple genes encoding for pro-inflammatory cytokines (e.g. *IL1B*, *IL6*, *IL12*, *IL23*) and chemokines (e.g. *CCL2*, *CXCL8*, *CXCL10*, *CXCL12*) in inflamed hiBMECs (Fig EV4B). Additionally, several genes related to antigen presentation through MHC-I and -II (e.g. *HLA-A*, *HLA-B*, *HLA-C*, *HLA-DRA*) were upregulated in inflamed hiBMECs, together with co-stimulatory molecules PD-L1 (*CD274*) and CD137 (*TNFRSF9*) (Fig EV4C).

To compare the extent of the inflammatory response elicited in hiBMECs versus hiECs, we performed differential gene expression analysis on inflamed hiBMECs versus inflamed hiECs. Notably, all genes mentioned above were downregulated in inflamed hiBMECs compared with inflamed hiECs (Fig 4C, Fig EV4D). We observed an overlap of 296 transcripts that were both upregulated in inflamed vs. control hiBMECs and downregulated in inflamed hiBMECs vs. hiECs (Fig 4D). In addition, GO enrichment analysis revealed an extensive overlap of over 400 terms associated with both these up- and downregulated genes (Fig 4E). Notably, the top 20 overlapping pathways were related with inflammatory processes and immune responses. These findings demonstrate that, while hiBMECs mount a robust inflammatory response to inflammatory stimuli, the magnitude of this response is significantly attenuated compared to naïve ECs, reflecting a more specialized and controlled inflammatory phenotype consistent with BBB ECs.

### HiBPCs and CHIR synergistically activate BBB pathways in hiBMECs

In our method, hiBMECs are differentiated from hiECs by the combined hiBPC co-culture and CHIR treatment. To dissect the contribution of each individual condition, we performed a principal component analysis, which revealed that 59% of total transcriptional variance can be explained by one principal component (PC1) splitting ECs treated with CHIR from untreated ECs (Fig 5A). Conversely, principal component 2 (PC2), accounting for 25% of total variance, distanced ECs co-cultured with hiBPCs from ECs in mono-culture.

**Figure 5.**
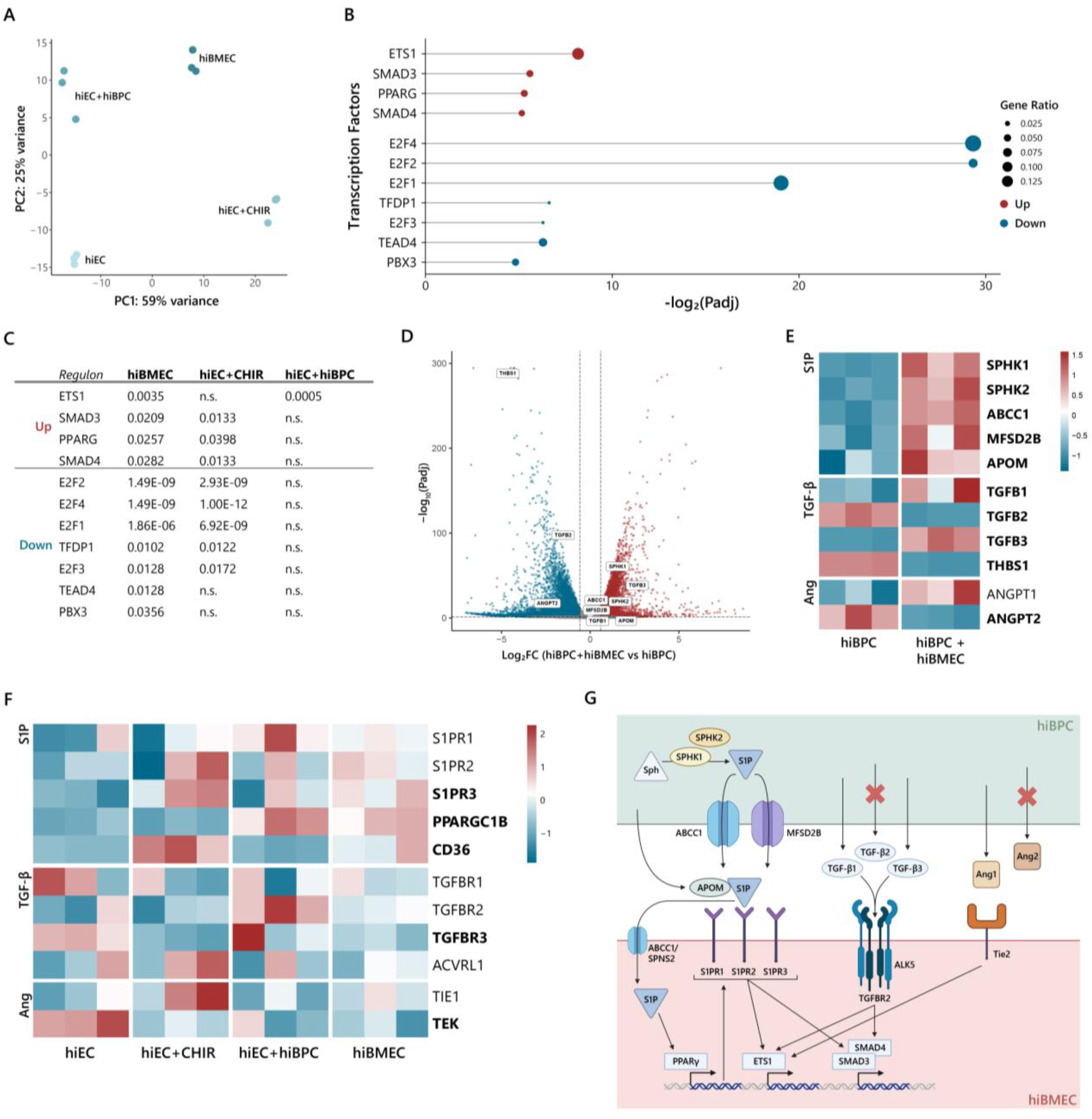
Synergistic activation of BBB-related transcriptional pathways in hiBMECs by hiBPCs and CHIR. **(A)** PCA plot of hiEC, hiEC+CHIR, hiEC+hiBPC and hiBMEC. **(B)** Transcription factor (TF) activity in hiBMEC vs hiEC, inferred using DoRothEA regulons. TFs with increased activity are marked in red, and TFs with decreased activity are marked in blue, ranked based on their adjusted *P* value (Padj). **(C)** Adjusted P values for the enrichment of DoRothEA regulon target genes among up- or downregulated genes in hiBMEC, hiEC+CHIR and hiEC+hiBPC compared to hiEC. **(D)** Volcano plot showing differentially expressed genes of hiBPC+hiBMEC vs. hiBPC. Upregulated genes (adjusted *p* value < 0.05 and fold change > 1.5) are displayed in red and downregulated genes (adjusted *p* value < 0.05 and fold change < −1.5) are displayed in blue. Transcripts related with pathways that activate the mentioned TFs are highlighted. N=3 per condition, GB4d hiPSC line. *P* values were determined using the Wald test and adjusted for multiple testing using the Benjamini-Hochberg procedure. **(E)** Z-score heatmaps of gene expression profiles across hiBPC and hiBPC+hiBMEC, and **(F)** hiEC, hiEC+CHIR, hiEC+hiBPC and hiBMEC. Selected transcripts are involved in S1P, TGF-β and Ang pathways. Significantly changed genes (hiBPC vs hiBPC+hiBMEC; hiEC vs hiBMEC) are marked in bold. **(G)** Schematic hypothetical overview of proposed signaling pathways regulating hiBMEC phenotype. Sphingosine (Sph) in hiBPC is phosphorylated by sphingosine kinases (SPHK1/2) to generate sphingosine-1-phosphate (S1P), which can be exported via ABCC1 and MFSD2B transporters. S1P, carried by apolipoprotein M (APOM), enters the hiBMEC cytoplasm or signals through S1P receptors (S1PR1-3), influencing downstream transcription factors including ETS1, SMAD3/4, and PPARγ. SMAD3/4 is also activated through binding of hiBPC-derived TGF-β1 and -β3, but not -β2, to ALK5/TGFBR2 in hiBMEC. Ang1, but not Ang2, is secreted by hiBPC and signals through the Tie2 receptor on hiBMEC to activate ETS1. Figure created in BioRender.

Differential gene expression analysis showed an upregulation of 1027 genes and a downregulation of 1328 genes in hiEC+CHIR versus hiECs. Of these, 398 were also upregulated and 647 downregulated in hiBMECs (Fig EV5A-B). Conversely, hiEC+hiBPC were enriched in only 239 genes and showed decreased expression of 254 genes when compared to hiEC, with 123 upregulated and 92 downregulated genes in common with hiBMECs. Notably, 281 and 396 genes were respectively up- or downregulated exclusively in hiBMECs and not in hiEC+hiBPC or hiEC+CHIR.

To further analyze the gene regulatory networks involved in hiBMEC differentiation, we assessed the differential activity of transcription factor (TF) regulons using DoRothEA, a resource for estimating TF activity from gene expression data. Out of eight repressed regulons in hiBMECs vs. hiECs, five were related to cell cycle (E2F2, E2F4, E2F1, TFDP1 and E2F3) (52, 53), two to angiogenesis (TEAD4 and PBX3) (54, 55) and one to endothelial inflammation (PBX3) (56) (Fig 5B). Moreover, out of 12 enriched regulons (Table S2), we identified four TFs that have been implicated in BBB establishment and regulation: ETS1, SMAD3, SMAD4 and PPARG (PPARγ). In particular, ETS1 stood out as the principal upregulated regulon in hiBMECs, with an adjusted P value of 0.0035 and a gene ratio of 0.08.

When analyzing the differential regulons in hiEC+CHIR and hiEC+hiBPC compared with hiEC, we observed that hiBPCs induce ETS1 activation, while CHIR induces most of the other regulons changed in hiBMECs (Fig 5C). Moreover, TEAD4 and PBX3 regulons are only significantly downregulated in hiBMECs, confirming the synergistic effects of hiBPC and CHIR on hiBMEC transcriptome.

To understand which molecular players drive these transcriptional changes, we interrogated whether paracrine ETS1 activators were differentially expressed in hiBPCs when co-cultured with hiBMECs. ETS1 pathway can be activated by multiple molecules, one of which being sphingosine-1-phosphate (S1P) (57). Notably, we observed an upregulation of the sphingosine kinases *SPHK1* (3.7 fold) and *SPHK2* (2.1 fold), the S1P transporters *ABCC1* (1.6 fold) and *MFSD2B* (2.3 fold), and the S1P carrier *APOM* (2.4 fold) in hiBPCs co-cultured with hiBMECs versus in mono-cultures (Fig 5D-E).

ETS1 can also be activated via transforming growth factor β (TGF-β) receptor signaling, through SMAD3/4 (58), whose regulons were also enriched in hiBMECs. Interestingly, hiBPCs co-cultured with hiBMECs showed an upregulation of *TGFB1* (1.2 fold) and *TGFB3* (5 fold) and downregulation of *TGFB2* (3.6 fold) and *THBS1* (19.8 fold), a major activator of latent TGF-β (59) (Fig 5D-E). Furthermore, angiopoietins are also able to activate the ETS1 pathway by binding to endothelial TEK (Tie2) (60). In accordance, hiBPCs in co-culture with hiBMECs strongly downregulated *ANGPT2* (6.7 fold), a Tie2 antagonist, and showed a trend towards upregulation of *ANGPT1*, a Tie2 agonist.

To dissect the intricate complexity of these transcriptional changes in hiBPCs, we further analyzed the individual effects of CHIR stimulation and hiEC co-culture on the expression of the genes mentioned above. We observed that most S1P-related genes (*SPHK1*, *SPHK2*, *ABCC1* and *MFSD2B*) were significantly upregulated by hiEC co-culture independently of the presence of CHIR (Fig EV5C). Moreover, hiECs induced a potent upregulation of *TGFB1* on hiBPCs, but CHIR dampened this effect, while also decreasing *TGFB2* and *THBS1* and increasing *TGFB3* expression. Conversely, hiBPCs co-cultured with hiEC downregulated *ANGPT1*. However, the addition of CHIR rescued this decrease and further induced the downregulation of *ANGPT2*.

On hiBMECs, CHIR increased the expression of *S1PR2*, *S1PR3* and the PPARγ target gene *CD36*, while hiBPCs induced the upregulation of *S1PR1* and the PPARγ co-activator *PPARGC1B* (Fig 5F). HiBPCs strongly upregulated *TGFBR2* in hiECs, but CHIR dampened this effect, while also downregulating *TGFBR3*, without significantly affecting the expression of other TGF-β receptors like ALK1 (*ACVRL1*) and ALK5 (*TGFBR1*).

Altogether, these results suggest a synergistic effect of hiBPCs and CHIR on regulating the transcriptomic changes that lead to the formation of hiBMECs from hiECs, converging on the ETS1, SMAD3/4 and PPARγ pathways (Fig 5G).

## Discussion

In this study, we describe a method to generate hiPSC-derived BMECs with robust barrier properties, transporter activity and adequate inflammatory responses. We showed, functionally and transcriptionally, that the synergistic effects of BPCs and Wnt/β-catenin pathway activation are necessary and sufficient to induce BBB properties in naïve endothelial cells. Moreover, this protocol was validated in three different hiPSC lines with three independent differentiations each, and in a physiologically relevant organ-on-chip platform, making it reproducible for future users across different *in vitro* systems.

HiBMECs exhibit barrier properties, characterized by the formation of TJs, achieving a TEER >80 Ω·cm² and NaFl permeability of 6×10⁻⁶ cm/s. These values represent an improvement compared to primary human BMECs, immortalized endothelial cell lines, and other hiPSC-derived BMEC-like cells (22, 61–63). While some protocols report TEER values >8000 Ω·cm², which are closer to estimated *in vivo* levels (1500–2000 Ω·cm²) (64, 65), these values are often derived from cells that also contain epithelial characteristics (UMM- and DMM-BMECs) (18, 19, 21), which have thicker morphology and more extensive TJ networks (66, 67). Consistent with the elevated TEER values, hiBMECs also show permeability to NaFl in the same order of magnitude of *in vivo* brain capillaries (68, 69). RNA sequencing further supports these findings, revealing the enrichment of multiple BBB markers in hiBMECs, such as *CLDN5*, *SLC2A1*, *SLC3A2*, and *ZIC3*, alongside downregulation of genes associated with transcytosis, including *PLVAP*, *CAV1*, and *CAVIN2*, which is indicative of the low vesicular trafficking typical of BMECs (72). Interestingly, *CLDN5* is significantly upregulated in hiBMEC and not in hiEC+CHIR or hiEC+hiBPC, demonstrating the synergistic effects of CHIR and hiBPCs in inducing the BMEC lineage in naïve ECs. Together, these findings highlight the suitability of hiBMECs for modeling key aspects of brain endothelial barrier function.

In addition to exhibiting proper barrier function, hiBMECs demonstrated functional expression of P-gp, a critical efflux transporter involved in physiological processes such as amyloid-β clearance and multi-drug resistance. However, not all studies on hiPSC-derived BMEC-like cells report on P-gp activity (20, 23). During human development, P-gp expression increases between gestation week (GW) 9 to 14 (44), coinciding with the emergence of pericyte-endothelial interactions around GW 11 (70, 71), while astrocyte-endothelial interactions happen only at GW 19 (72). In line with this, we observed P-gp activity in hiECs co-cultured with hiBPCs, but not with hiPSC-derived astrocytes or hiBPC conditioned medium. Given its dimensions and morphology, the structures that physically connected hiBPCs and hiBMECs on inserts appear to be tunneling nanotubes (73, 74), through which biomolecular cargo can be transferred directly from the hiBPC to the hiBMEC cytoplasm and modulate P-gp function independently of paracrine signaling. Furthermore, we observed that hiBPC co-culture induces a strong downregulation of *EDN1* in hiBMECs. Endothelin-1 is a vasoconstrictor that inhibits P-gp activity through phosphorylation of caveolin-1 (75–77). Thus, its downregulation might explain the induction of P-gp activity observed in hiBMECs. Together, our results highlight the importance of direct pericyte-endothelial interactions in establishing and maintaining functional efflux properties in hiBMECs.

It is known that BMECs have a low expression of CAMs during homeostatic conditions, and their inflammatory response is tightly regulated to maintain the integrity of the BBB and protect the brain from immune cell infiltration, contrary to peripheral ECs (78). Yet, to the best of our knowledge, no study has directly compared the effects of inflammatory stimuli on BMECs and peripheral ECs. Here, we found that hiBMECs responded to inflammatory stimuli by upregulating CAMs, cytokines, and antigen presentation molecules, but less pronouncedly than hiECs. Particularly, we observed that 90% of inflamed hiBMECs were positive for ICAM-1, which is in line with a seminal study showing that more than 80% of vessels in MS lesions are ICAM-1^+^, in contrast with 10% in non-neurological control brains (79). Conversely, only 9% of inflamed hiBMECs were positive for VCAM-1, which also aligns with previous work indicating that aging and exposure to systemic inflammatory mediators increases the percentage of VCAM-1^+^ brain vessels from 1-3% to 5-10% in mice (80). Overall, despite the lower VCAM-1 upregulation in inflamed hiBMECs compared to other protocols (20, 81), the percentages we observed across donors indicate an appropriate response of hiBMECs to inflammation.

To better understand the molecular mechanisms underlying BBB induction in hiBMECs, we first benchmarked our RNA sequencing dataset against publicly available datasets. Unlike other hiPSC-derived BMEC-like cells, hiBMECs clustered closely with primary and freshly isolated adult BMECs, suggesting a more physiologically relevant transcriptional profile. An important upregulated regulon in our dataset is ETS1, a TF that is highly expressed in BMECs and promotes vascular integrity while suppressing endothelial-to-mesenchymal transition (82, 83). We pinpointed three pathways that may explain the BBB induction via ETS1 in hiBMECs, potentially triggered by inducers secreted by hiBPCs observed in our study, namely Ang1/Tie2, S1P and TGF-β/ALK5 (Fig 5G). (84). Angiopoietins are important modulators of vascular and BBB development. Ang1 secreted by pericytes stabilizes the BBB by activating Tie2 receptors on endothelial cells (85, 86). Conversely, angiopoietin-2 (Ang2) antagonizes Tie2 signaling, disrupting junctional proteins, increasing vesicular permeability, and causing pericyte detachment (87–89). We observed that CHIR induces a strong upregulation of *ANGPT1* and downregulation of *ANGPT2* on hiBPCs, which may explain some of the properties attained by hiBMECs upon differentiation.

Another ETS1 activator described in literature is the lipid mediator S1P (57), which is a potent BBB modulator that stabilizes the cytoskeleton and enhances cell-to-cell contact through recruitment of adherens and tight junctions to the cell membrane, as we demonstrated before (90, 91). Interestingly, we observed an upregulation of the sphingosine synthesis and transport machinery *SPHK1*, *SPHK2*, *ABCC1*, *MFSD2B* and *APOM* in hiBPCs co-cultured with hiBMECs, suggesting that hiBMECs induce the production and facilitate the transport of S1P from hiBPCs. This is in line with a previous report indicating that pericyte-derived S1P maintains barrier properties in retinal ECs (92). However, the role of S1P at the BBB is not limited to its interaction with ETS1. On the one hand, S1P can enter hiBMECs through ABCC1 and SPNS2 and bind to PPARγ, activating it (93). PPARγ, another regulon enriched in hiBMECs, enhances endothelial barrier function by reducing inflammation and oxidative stress (94), while inducing S1PR1 expression (95). On the other hand, S1P can bind to S1PRs, which are also upregulated in hiBMECs, activating not only ETS1 but also SMAD3 and SMAD4 (96, 97), two additional regulons enriched in hiBMECs.

SMAD3 acts in concert with ETS1 to stabilize newly formed microvessels (98), while SMAD4 stabilizes BMEC-BPC interactions (99). Both SMAD3 and 4 are involved in the TGF-β/ALK5 signaling pathway, which has been shown to inhibit angiogenesis and enhance vascular stability (100). In contrast, signaling through ALK1 and SMAD1/5 induces EC migration and angiogenesis (100, 101). TGF-β1 strongly activates both ALK1 and ALK5; however, at high concentrations, ALK1 antagonizes ALK5 (102). Therefore, TGF-β1 drives a biphasic response in ECs where low concentrations activate ALK5, promoting vessel stability, while high concentrations activate ALK1, inducing angiogenesis (100). In this study, we observed a potent upregulation of *TGFB1* in hiBPCs co-cultured with hiECs, which was dampened by CHIR stimulation. Although previous studies have described improved endothelial barrier properties upon TGF-β pathway inhibition (15, 35, 103), considering the biphasic response of ECs to TGF-β1, we postulate that this balanced *TGFB1* upregulation in hiBPCs leads to activation of the stabilizing ALK5/SMAD3 pathway on hiBMECs. Altogether, these results suggest a reciprocal interaction between hiECs and hiBPCs, where hiECs drive hiBPCs to a supportive BBB-inducing phenotype, which in turn activate multiple signaling pathways that induce BMEC commitment in hiECs.

In conclusion, we showed in this study that BPCs and Wnt/β-catenin signaling act in concert to induce BMEC lineage commitment in human naïve endothelium. The presence of key BBB characteristics in our hiBMEC-hiBPC model makes it suitable for future toxicity screening, testing of novel drug delivery vehicles, and disease modeling, including cerebrovascular and neurodegenerative disorders such as stroke, Alzheimer’s disease, and multiple sclerosis. Additionally, further studies should focus on validating the proposed BBB-inducing pathways at a functional level, for fundamental and therapeutical purposes.

## Materials & Methods

### Donors

GB2a (RRID: CVCL_D0PU) and GB4d (RRID: CVCL_D0PT) lines were derived from dermal fibroblasts obtained from Tissue Solutions. The reprogramming of these lines were exempt from ethical approval, as decided by the Amsterdam UMC Medisch Ethische Toetsings Commissie (METC). TMO line (RRID: CVCL_RM92) was purchased from Gibco. SCVI111 (RRID:CVCL_C6U9) was obtained from Stanford Cardiovascular Institute.

### Reprogramming and culture of hiPSCs

GB2a and GB4d fibroblasts at passage 5 were dissociated into single-cell suspensions using 0.05% Trypsin/EDTA (Gibco). For each cell line, 7.5 × 10⁵ cells were resuspended in 82 µL nucleofection buffer combined with 18 µL supplement from the P2 Primary Cell 4D-Nucleofector X kit (Lonza), along with 1.67 µL each of Addgene plasmids #27080, #27078, and #27076 (2 µg/µL) (104). The cell-plasmid mixture was transferred to a nucleofection cuvette and electroporated using an Amaxa 4D-Nucleofector (Lonza) with program DT-130. Immediately post-nucleofection, 500 µL of penicillin/streptomycin-free fibroblast medium [DMEM (Gibco) + 1% Ultroser G (Sartorius)] was added dropwise to the suspension, followed by a 10-minute incubation. The cells were then plated onto a 10-cm culture dish containing 10 mL of the same medium and incubated overnight at 37°C with 5% CO₂. After 48 hours, the medium was replaced with fresh fibroblast medium containing penicillin/streptomycin. Seven days post-nucleofection, cells were washed with phosphate buffer saline (PBS), detached using Trypsin/EDTA, collected in fibroblast medium, and centrifuged at 300g for 5 minutes. The pellet was resuspended in fresh fibroblast medium and passaged at a 1:3 ratio onto 10-cm dishes coated with LN521 (Biolamina). After 24 hours, the medium was switched to TeSR-E7 (STEMCELL Technologies), and 24 hours later cells were transferred to a 5% O₂ incubator to enhance reprogramming efficiency. Medium was refreshed daily until hiPSC colonies appeared (3–4 weeks). Colonies were manually dissected using insulin needles (Becton Dickinson) and replated onto LN521-coated wells in mTeSR Plus medium. Cells were passaged ≥10 times, and pluripotency was verified using the STEMdiff Trilineage Differentiation Assay Kit (STEMCELL Technologies) and immunocytochemistry for mesodermal (Brachyury), endodermal (SOX17, AFP), and ectodermal (β3-tubulin, PAX6) markers. Post-passage 10, hiPSCs were transitioned to 20% O₂ and cultured on vitronectin-coated surfaces. All lines were maintained by refreshing mTeSR Plus every 48 hours and passaged as colonies at 70–80% confluence using Versene (Gibco).

### Differentiation of hiBPCs

GB2a, GB4d, and TMO hiPSC lines were differentiated into neural crest (NC)-derived brain pericytes (hiBPC) using an adaptation of previously published protocols (105, 106). Briefly, hiPSCs were passaged as single cells and seeded at a density of 2×10^5^ cells/cm^2^ onto Vitronectin-coated plates. Cells were cultured in NC induction medium, consisting of DMEM/F12 GlutaMAX (Gibco), 1x B27 (Gibco), 0.5% BSA and 3 µM CHIR 99021 (CHIR; Tocris) for 5 days, with daily media refreshments. 10 μM ROCK-inhibitor (Y-27632; Selleck Biotechnology) was added on the first day. The resulting NC cells were singularized with Accutase (Innovative Cell Technologies) and frozen in CryoStor CS10 (STEMCELL Technologies), or directly seeded onto 0.1% gelatin-coated plates at a density of 2.5 ×10^4^ cells/cm^2^. NC cells were cultured in pericyte medium (PM; ScienCell) for 5 days for BPC specification, with daily refreshments. Immunocytochemistry and mRNA evaluation of PDGFRβ, NG2, CD13, CD146, FOXF2 and FOXC1 confirmed their pericyte identity. HiBPC were used between passages 3 and 6.

### Differentiation of hiECs

GB2a, GB4d, TMO and SCVI111 hiPSC lines were differentiated into naïve endothelial cells using an adaptation of previously published protocols (30, 31). Briefly, hiPSCs with a confluency of 70 – 80% were detached using Versene and pipetted up-and-down to obtain clumps with the size of 0.5 – 1 mm. hiPSC clumps were seeded onto vitronectin-coated plates or flasks at a passage ratio of 1:20 – 1:30, depending on the confluency and growth speed of the hiPSC line. hiPSCs were cultured for 24h in mTeSR Plus. At day 0 of differentiation, mesoderm induction was started by changing the medium to B(P)EL with 8 μM CHIR. At day 3 of differentiation, vascular specification was started by changing the medium to B(P)EL with 50 ng/mL vascular endothelial growth factor (VEGF; Biolegend) and 10 μM SB431542 (Selleck Biotechnology). Medium was refreshed on days 6 and 8. At day 10 of differentiation, cells were detached using Accutase, centrifuged at 300g for 5 minutes, counted, and magnetically-activated cell sorted for PECAM-1 using the CD31 MicroBead Kit (Miltenyi Biotec), the QuadroMACS™ Separator (Miltenyi Biotec) and the LS columns (Miltenyi Biotec), following the manufacturer’s instructions. PECAM-1^+^ endothelial cells (hiECs) were counted and frozen in CryoStor CS10, or seeded at a density of 1.5×10^4^ cells/cm^2^ onto plates or flasks coated with 10 ug/mL human placenta-derived collagen IV (Sigma-Aldrich) in endothelial serum-reduced medium ++ (ECSR++), comprising of human endothelial serum-free medium (Gibco), 1% human serum derived from platelet-poor plasma (Sigma-Aldrich), 30 ng/mL VEGF and 20 ng/mL basic fibroblast growth factor (bFGF; Peprotech). HiECs were cultured in ECSR++ and passaged up to 3 times at 1:4 ratio when 100% confluency was reached.

### Differentiation of hiBMECs on inserts

The apical side of 12-well or 24-well transparent polyethylenterephthalate (PET) cell culture inserts with 1 µm pore size (cellQART) was coated with 10 ug/mL collagen IV and the basolateral side of the same inserts was coated with 0.1% gelatin. After 2 hours, coatings were aspirated and plates were inverted to access the basolateral side of the inserts. HiBPCs were detached with Accutase, centrifuged at 300g for 5 min, counted, resuspended in PM, and plated at 18.000 cells/cm^2^ on the basolateral side of the insert in a droplet. Cells were incubated for 1h at 37°C for attachment. HiECs were detached with Accutase, centrifuged at 300g for 5 min, counted, resuspended in ECSR++, and plated at 60.000 cells/cm^2^ on the apical side of the insert. PM medium was added to the bottom well. Cell concentrations and volumes can be found on table 1. Cells were cultured for 6 days, with media changes every two to three days. On day 4 of co-culture, 4 μM CHIR was added to the apical side of the inserts.

**Table 1.** Cell concentrations and volumes used for 12-well and 24-well inserts.

|  | 12-well insert (1.1 cm<sup>2</sup>) | 24-well insert (0.3 cm<sup>2</sup>) |
| --- | --- | --- |
| **Step 1** |  |  |
| hiBPC concentration | 200.000 cells/mL | 100.000 cells/mL |
| hiBPC volume | 100 μL | 60 μL |
| **Step 2** |  |  |
| hiEC concentration | 130.000 cells/mL | 90.000 cells/mL |
| hiEC volume | 500 μL | 200 μL |
| **Step 3** |  |  |
| PM volume | 1400 μL | 740 μL |

To induce inflammation, ECSR++ with 25 IU/mL TNFα (Peprotech) and 25 IU/mL IFNγ (Peprotech) were added to the apical side of the inserts and incubated for 16 hours at 37°C.

### Differentiation of hiPSC-derived astrocytes

GB4d hiPSC line was differentiated into astrocytes using an adaptation of a previously published protocol (107). Briefly, hiPSCs were washed twice with 0.05% EDTA for approximately 30 s and left in PBS until colonies came loose by tapping the plate (5 to 10 min). The fragments were collected in 5 mL PBS and left in a tube until the fragments deposited. The PBS was aspirated and the fragments were resuspended in 3 mL Neuro Maintenance Medium (NMM; 1:1 DMEM + GlutaMAX (Gibco) : Neurobasal Medium (Gibco), 1x B27, 1x N2 (Gibco), 2.5 μg/mL insulin (Sigma-Aldrich), 1.5 mM L-glutamine (Gibco), 100 μM non-essential aminoacids (Gibco), 50 μM 2-mercaptoethanol, 1% penicillin/streptomycin) containing 20 ng/mL epidermal growth factor (EGF; Peprotech), 4 ng/mL bFGF, 40 ng/mL T3 (Sigma-Aldrich) and 10 μM Y-27632, and transferred to an anti-adhesive poly 2-hydroxyethyl methacrylate-coated plate. Embryoid bodies formed overnight and were plated on day 3 onto Geltrex (Gibco)-coated plates in 3 mL NMM containing 20 ng/mL EGF, 4 ng/mL bFGF, 40 ng/ml T3, 10 μM RA (Sigma-Aldrich) and 10 μM Y-27632. Two-third medium changes were performed every other day until day 10. At day 10, medium was changed into NMM with 20 ng/mL EGF and 40 ng/mL T3 and two-third of the medium was changed every other day until day 18. Cells were passaged with Accutase when confluent. At day 18, medium was changed into NMM without vitamin A with 20 ng/mL EGF and 40 ng/mL T3. Two-third of the medium was changed every other day until day 42, when medium was changed into Astrocyte Medium (ScienCell). Thereafter, half medium changes were done every other day and cells were passaged at confluence using Accutase. Immunocytochemistry or mRNA evaluation of GFAP, S100B, ALDH1L1, GLAST, Vimentin, AQP4 and Kir4.1 confirmed their astrocyte identity. Astrocytes were used between day 80 and 100.

### Immunocytochemistry on inserts

Cells were carefully rinsed with phosphate-buffered saline (PBS) once and fixed with ice-cold methanol or 4% paraformaldehyde (PFA) for 10 min. In case of PFA fixation, cells were further permeabilized for 10 min using 0.3% Triton-×100 in PBS (Sigma-Aldrich). Cells were then blocked with 10% goat serum in 0.05% Tween 20 (Sigma-Aldrich) in PBS (PBSGT) for 30 min. Primary antibodies were diluted in 1% PBSGT, and cells were incubated in the antibody solutions at 4°C overnight or at room temperature for 2 hours. After three PBS washes, cells were incubated with secondary antibodies and DAPI in 1% PBSGT for 2 hours at room temperature. Cells were then washed with PBS three times, and fluorescence imaging was performed using the Nikon AX R or the Leica SP8 confocal microscopes. A detailed list of antibodies is shown in table S3.

### Scanning electron microscopy

Inserts with hiBMECs and hiBPCs were fixed using pre-warmed (37°C) 4% EM-grade paraformaldehyde and 1% EM-grade glutaraldehyde (EMS) in 0.1 M phosphate buffer for 4 hours at room temperature. The fixation procedure was performed promptly to prevent the samples from drying before fixative application. Following the 4-hour fixation, samples were stored at 4°C until further processing. For dehydration, samples were washed three times with water for 5 minutes each. They were then dehydrated with increasing concentrations of ethanol: 30%, 50%, 70%, 80%, 90%, 96%, and twice in 100%, each for 10 minutes. After completing the dehydration series, samples were placed in a critical point dryer (CPD300, Leica) using a 16-cycle protocol at slow speed. Membranes were removed from the inserts and cut into approximately 1 mm strips, which were positioned upright on scanning electron microscopy stubs with the cut edge facing up. The stubs were sputter-coated with palladium/platinum using a sputter coater (ACE 600, Leica). Images were acquired with a Zeiss Gemini 300 Sigma scanning electron microscope.

### Transendothelial electrical resistance measurement

TEER was measured using an EVOM3 voltohmmeter (World Precision Instruments) equipped with a STX4 chopstick electrode (World Precision Instruments) on 12-well inserts at day 16. Electrodes were sterilized with 70% ethanol and equilibrated in pre-warmed PBS for 5 min. TEER in Ω·cm^2^ was calculated by subtracting the resistance of one coated cell-free insert from resistance values of each sample, multiplied by the surface area of the insert.

### Permeability assay on inserts

10 μM sodium fluorescein (NaFl, Sigma-Aldrich) was diluted in ECSR++ and added to the apical side of the inserts. Cells were incubated for 1 hour at 37°C and a calibration curve of serial dilutions of NaFl was prepared in a black wall 96-well plate. Samples from the basolateral compartment were collected and the fluorescence was measured using a Synergy HTX Multimode Reader (Biotek), with excitation/emission wavelengths of 485/530 nm. Apparent permeability coefficients (P_app_) were calculated using the following equation: P_app_ = (C_b_ × V_b_)/(C_a_ × A × Δt), where C_b_ is the concentration measured in the basolateral compartment at 1 h, V_b_ is the volume of the basolateral compartment (0.8 mL for 24-well inserts), C_a_ is the concentration in the apical compartment at 0 h (10 μM), A is the surface area available for permeability (0.3 cm^2^ for 24-well inserts) and Δt is the duration of the experiment (3600 seconds).

### P-glycoprotein activity assay on inserts

The P-gp activity was evaluated by assessing the permeability to rhodamine 123 (Rho_123_), a fluorescent P-gp substrate, with or without the presence of verapamil, a P-gp blocker. The respective groups were pre-incubated with 25 μM verapamil (Sigma-Aldrich) diluted in ECSR++ and added to the apical side of the inserts 1 hour prior to Rho_123_ treatment. After 1 hour, 10 μM Rho_123_ (Sigma-Aldrich) diluted in ECSR++ was added to the apical side of the inserts and cells were incubated for 1 hour at 37°C. Samples from the basolateral compartment were collected in a black wall 96-well plate and the fluorescence was measured using a Synergy HTX Multimode Reader, with excitation/emission wavelengths of 485/530 nm. Concentrations of Rho_123_ were determined by standard calibration curves. P_app_ coefficients were calculated as described above. P-gp activity coefficient was calculated using the equation (P_app_^VP^/P_app_^UNT^)-1, where P_app_^VP^ is the Rho_123_ P_app_ on VP-treated cells and P_app_^UNT^ is the Rho_123_ P_app_ on untreated counterparts.

### Flow cytometry

Cells were dissociated with TrypLE Select (Gibco), centrifuged at 300g for 5 min, and suspended in flow buffer (PBS + 0.1% BSA). Cells were transferred to 96-well V-bottom plates, centrifuged (1500 rpm, 5 min), and stained with viability dye in PBS for 15 min at room temperature. After centrifugation (1500 rpm, 3 min), cells were washed with 100 µL flow buffer. Surface antibodies (50 µL/well) were added, and cells were incubated for 30 min at 4°C. Cells were washed three times with flow buffer (centrifuging at 1500 rpm, 3 min each), then suspended in flow buffer and transferred to round-bottom tubes. Acquisition was performed on a BD LSRFortessa cytometer (Becton Dickinson) and analyzed using FlowJo. A detailed list of antibodies is shown in table S3.

### Generation of BBB-on-chip (BBBoC)

BBBoC generation was adapted from previously published protocols (108, 109) with minor adjustments. In brief, commercially available microfluidic chips idenTx 9 (AIM Biotech) with one gel channel and two flanking media channels were used. To cast BBBoCs, SCVI111 hiECs and GB4d hiBPCs were brought into suspension at a concentration of 10 × 10^6^ hiECs/mL and 3.3 × 10^6^ hiBPCs/mL to obtain a cell seeding ratio of 3:1. Cells were resuspended in ECGM-2 (Promocell) supplemented with 50 ng/mL VEGF and 4 U/mL bovine thrombin (Sigma-Aldrich). Prior to seeding, 1 part cell mix was resuspended in 1 part ECGM-2 with 50 ng/mL VEGF and 4 U/mL thrombin, which was further mixed in a 1:1 ratio with 6 mg/mL human fibrinogen (Sigma-Aldrich) for a final concentration of 3 mg/mL. To cast the cell/hydrogel mix in the microfluidic chip, 15 µL of cell/hydrogel mix was injected into the middle chamber of the idenTx 9 platform and incubated at room temperature for 15 minutes to allow enzymatic crosslinking. After this, side channels were hydrated with 15 mL of ECGM-2 with VEGF and gravity-driven flow was induced by adding 100 µL ECGM-2 with VEGF to the right media inlets and 50 µL to the left media inlets. Medium was replaced daily in a respective manner to maintain gravity driven flow over 7 days. On day 1, all chips were supplemented with the γ-secretase inhibitor DAPT (10 μM, Tocris) to promote hiEC sprouting, and on day 6 and 7 CHIR-treated chips were supplemented with CHIR to complete hiBMEC differentiation.

### Immunocytochemistry on BBBoCs

BBBoCs were fixed directly in the microfluidic devices using 4% PFA or ice-cold methanol for 30 minutes at room temperature. Permeabilization of cells was enabled by 0.5% Triton X-100 treatment for 15 minutes at room temperature. Each step was followed by three washes with PBS for 10 minutes each, and samples were subsequently blocked in PBS containing 2% bovine serum albumin (BSA; Sigma-Aldrich) for 3 hours at room temperature. Primary antibodies were diluted 1:400 in 1% BSA and incubated overnight at 4 °C. Antibodies used are listed in table S3. After primary antibody incubation, samples were washed three times with PBS for 15 minutes, followed by incubation with secondary antibodies (diluted 1:600 in 1% BSA) for 2 hours at room temperature. Fluorescence imaging of entire microfluidic channels was performed using a Nikon AX R confocal. Image processing and 3D reconstruction were conducted using Imaris software (version 10.2.0, Oxford Instruments), NIS-Elements software or ImageJ (National Institutes of Health, Bethesda, MD, USA), depending on the quantification.

### Geometry analysis of vascular network in BBBoC

To assess vascular morphology, networks were fixed at day 7 and immunostained (as described above) using either DyLight 649-labeled Ulex Europaeus Agglutinin I (UEA I; Vector Laboratories) or PECAM-1 antibody to visualize vessel structures. Thresholding parameters for binarization were determined based on representative overview images to ensure consistent segmentation of vascular networks across samples. Image binarization was performed using CellProfiler (version 4.2.1, Broad Institute). Binarized images were analyzed using the DiameterJ plugin for ImageJ (110), which applies automated skeletonization and morphometric analysis to quantify vascular network geometry. Measured parameters included average diameter, average branch length, total network length and number of branching points. All image processing and analysis steps were applied uniformly to maintain consistency across experimental conditions.

### Permeability assay on BBBoCs

After seven days of culture, a permeability assay was performed. To visualize vascular networks, BBBoCs were incubated with UEA I for 1h in a CO₂ incubator. Live-cell fluorescence imaging was performed using a Nikon AX R confocal microscope equipped with environmental control (37°C, 5% CO₂). 70-kDa Tetramethylrhodamine isothiocyanate-dextran (TRITC-dextran, Sigma-Aldrich, 2 µg/mL) in ECGM-2 medium was added into the top left (100 µL) and right (50 µL) inlets of the BBBoCs. Images were acquired every 10 minutes over a 50-minute period. Quantification was performed on NIS-Elements software (version 5.42.04, Nikon Instruments Inc.). In brief, regions of interest (ROIs) were selected based on UEA I^+^ (vessel) and UEA I ^−^ (gel) threshold areas was performed. Subsequently, the MFI of TRITC-Dextran in each ROI was averaged and used to quantify the changes in fluorescence in the gel over time.

### P-glycoprotein activity assay on BBBoC

After seven days of culture, the P-gp activity was evaluated using Rho_123_ and verapamil. The medium of all BBBoCs was replaced with ECGM-2 with UEA I (1:600) and with or without 25 μM verapamil 1 hour prior to Rho_123_ treatment. After 1 hour, media were aspirated and 10 μM Rho_123_ diluted in ECGM-2 was added into the top left (100 µL) and right (50 µL) inlets. After 1 hour at 37°C, BBBoCs were moved to a Nikon AX R confocal microscope equipped with environmental control (37°C, 5% CO₂) for live-cell fluorescence imaging, and images were captured at 0 and 1 hours of imaging. Quantification was performed on NIS-Elements software in a similar manner to the BBBoC permeability assay. Briefly, ROIs were selected based on UEA I^+^ (vessel) and UEA I^−^ (gel) areas. Subsequently, the MFI of Rho_123_ in each ROI was averaged and used to quantify the changes in fluorescence in the gel and the vessel on both time points.

### RNA sequencing and data analysis

Total RNA from cells was isolated in TRIzol (Invitrogen) and a TapeStation RNA ScreenTape assay (Agilent) was performed to assess RNA integrity. RNA libraries were prepared and multiplexed using KAPA mRNA Hyperprep (Roche), and used as the input for high-throughput sequencing via NovaSeq X Plus platform (Illumina). Previously published RNA sequencing data of hiPSC-derived BMEC-like cells and various EC and epithelial cell types were downloaded from the Gene Expression Omnibus (GEO accession numbers: GSE157852, GSE57662, GSE173206, GSE224846, GSE243193, GSE97575, GSE108012, GSE129290, GSE256493, GSE137619, GSE271935) and processed the same way as the libraries prepared in this study.

Raw RNA sequencing data were first assessed for quality using FastQC (111) and then aligned to the human genome using STAR (version 2.7.10a) (112). Genome indices were generated from the reference FASTA and GTF annotation files prior to alignment. Post-alignment, the resulting SAM files were converted to BAM format, sorted, and indexed using Samtools (version 1.13) (113). Gene-level read quantification was performed using featureCounts (version 2.0.3) (114) from the Subread package (115), employing the corresponding GTF annotation file. For differential gene expression analysis, raw count data were imported into R (version 4.4.1) and analyzed using the DESeq2 package (version 1.44.0) (116). Genes with less than 5 counts in at least 3 different samples were filtered prior to normalization. Differential gene expression was determined using the Wald test, and p-values were adjusted for multiple testing using the Benjamini-Hochberg procedure. Genes with an adjusted p-value < 0.05 and |log_2_ fold change| > 1.5 were considered significantly differentially expressed. Heatmaps were created using the pheatmap package with normalized z-scores and applying gene clustering. Volcano plots using were created using the EnchancedVolcano package.

Gene ontology (GO) enrichment analysis was performed using ShinyGO (v0.80) to identify overrepresented Biological Process (BP) categories among the differentially expressed genes (117). The input gene list consisted of significant differentially expressed genes with an adjusted p-value < 0.05 and |log₂ fold change| > 1.5 from the DESeq2 analysis. Enrichment was assessed based on a hypergeometric test with false discovery rate (FDR) correction for multiple testing. GO terms with FDR-adjusted p-values < 0.05 were considered significantly enriched. Selected enriched terms were visualized using lollipop plots created in SRplot (118).

### Statistical analysis

Data were statistically analyzed using GraphPad Prism v10 (GraphPad Software). Normal distribution was determined using Shapiro-Wilk test. All datasets were normally distributed, thus parametric tests were used. For datasets with 2 groups, two-tailed paired student t-test was used. For datasets with 3 or more groups, ordinary one-way ANOVA was used. For datasets with 2 groups and measurements over time, two-way repeated measures ANOVA with Greenhouse-Geisser correction was used. *P < 0.05, **P < 0.01, ***P < 0.001.

## Data availability

RNA sequencing data generated and analyzed in this study have been deposited to the Gene Expression Omnibus (GEO) repository under accession number GSE305015.

## Acknowledgements

The authors acknowledge gratefully the Microscopy and Cytometry Core Facility (MCCF) of Amsterdam UMC for their expertise and support in microscopy and flow cytometry, the Core Facility Genomics (CFG) of Amsterdam UMC for the performance of RNA sequencing, and the Cellular Imaging Facility of Amsterdam UMC for their expertise and support in scanning electron microscopy.

This work was funded by CONNECT project (NWO Human Measurement Models 2.0, project number: 18957), Horizon ERC Advanced (ERC-AD 2022: 101097983), and IMM-BBB project (co-funded by PPP Subsidy awarded by Health∼Holland, Top Sector Life Sciences & Health, in collaboration with the BRAINS program) to HEdV. NMdW was funded by ZonMw Onderzoeksprogramma Dementie (project number: 10510022110005). JPNB, SDBH and SG were funded by Body Barriers project (NWO AES programme; project number: 19247). PCH and PLH were funded by Amsterdam UMC (project number: 3910).

## Author contributions

HNP performed experiments, analyzed data, designed and conceived the study and wrote the manuscript. NRK performed experiments, contributed to design the study and experiments, and revised the manuscript. PCH performed and designed experiments, analyzed data, and revised the manuscript. MKvD and SMAvdP performed experiments. MdK provided support on RNAseq analysis. NJP performed the scanning electron microscopy imaging and sample preparation. JPNB performed the characterization and quality control of hiPSC lines generated in-house. PLH and SG provided material and valuable scientific input, and revised the manuscript. SDBH generated hiPSC lines, supervised the study, and contributed to design the study. NMdW and HEdV supervised the study, contributed to design the study, obtained funding and revised the manuscript. All authors read and approved the publication of this manuscript.

## Disclosure and competing interests statement

The authors declare no competing interests.

## Notes

### Competing Interest Statement

The authors have declared no competing interest.

